# Aging alters mRNA processing in the mouse ovary

**DOI:** 10.1101/2025.04.16.649248

**Authors:** Kevin Vo, Grace J. Pei, Ramkumar Thyagarajan, Patrick E. Fields, M. A. Karim Rumi

**Affiliations:** Pathology and Laboratory Medicine, University of Kansas Medical Center, Kansas City, KS 66160, USA; Internal Medicine (GERI), University of Kansas Medical Center, Kansas City, KS 66160, USA

**Keywords:** Ovary, aging, gene expression, pre-mRNA processing, transcript variants, RNA binding proteins

## Abstract

Aging in females predominantly impacts the ovaries before any other organ systems. This has profound implications for women’s reproductive health. This phenomenon is closely linked to a gradual depletion of the ovarian follicle reserve and a notable diminishment of oocyte quality. Studies have shown that cellular changes within ovaries can manifest even before the observable depletion of ovarian follicles. To understand the molecular mechanisms underlying these changes, we have conducted a comprehensive analysis of the changes in gene expression in aging mouse ovaries. Using efficient genomics software such as CLC Genomic Workbench, we could detect not only the differentially expressed genes but also delineate the various transcript variants present in the transcriptome of aging ovaries. We verified the results by comparing coding sequences of selected transcripts with the coding sequences of their canonical counterparts from young and aged mice. In general, the analysis methods yielded similar observations. Our findings revealed that traditional gene expression analyses often overlook the differential expression of numerous transcript variants. We identified significant alterations in the expression patterns of alternative transcripts in aging ovaries and found coding sequences that lead to profound functional outcomes. Notably, most of these differentially expressed transcript variants were affected upstream epigenetically and transcriptionally, then generated through alternative splicing events. This suggests that aging may lead to alterations in RNA-binding proteins and spliceosome components, which play a crucial role in mRNA processing within the mouse ovary. Our observations highlight the necessity of focusing on transcript variants and their functions in aging research, as they provide a more nuanced understanding of the biological processes at play.

## 1. Introduction

Ovarian aging is a poorly understood natural process of declining ovarian functions: steroidogenesis and oogenesis [1]. This natural process impacts all women after 45 years, which remains a significant hurdle to general health. An age-induced deficit in the hypothalamic-pituitary axis and changes within the ovary occur during ovarian aging [2,3]. Several factors accelerate ovarian aging, such as oxidative stress, cohesion deterioration, cellular senescence, gene mutations, autoimmunity, chemotherapy, and radiotherapy [1,4-6]. These pathological conditions can result in premature ovarian insufficiency (POI) in younger women, which differs from ovarian aging [7,8]. The decline in ovarian follicle reserve is the primary cause of ovarian aging [9].

Changes in cellular functions depend on gene expression, which is regulated at epigenetic, transcriptional, post-transcriptional, translational, and post-translational levels [10,11]. Epigenetic and transcriptional regulations are intricately linked processes and represent dynamic changes in the cellular microenvironment [12]. These molecular mechanisms determine the quality and quantity of mRNA in a cell.

Epigenetic regulation affects chromatin structure and transcription factor binding, allowing for the enhancement or repression of transcription [13]. One of the main modifications caused by epigenetics includes DNA methylation [14]. Consequently, reproductive aging undergoes dynamic alterations in methylation, causing high levels of methylation in mature oocytes [15]. These high DNA methylation levels regulate gene expression, leading to tissue-specific functions, and play a role in aging. Over time, these oocytes decrease in quantity and may reflect a decrease in DNA methylation or an increase in demethylation with aging oocytes[16]. Transcription factors code for specific proteins that regulate gene expression through DNA sequence binding [17]. Although only a small portion of transcription factors are seen to be involved in ovarian functions, the effects of such factors can be highly detrimental to ovarian development [18]. The forkhead transcription factor, *FOXL2*, is required for granulosa cell functions such as differentiation and proliferation [18]. A repression of the *FOXL2* factor will impair the number of functional granulosa cells and eventually lead to oocyte atresia and infertility [19]. *FOXL2* is not a direct cause of ovarian aging, but it demonstrates the effects of transcriptional regulation on the process.

Outside of epigenetic and transcript regulation, RNA-binding proteins can also regulate gene expression [20]. They can dictate post-transcriptional processes, such as pre-mRNA splicing [21]. Post-transcriptional modifications of RNA are essential for both pre-mRNA processing and mRNA functions to generate mature mRNAs and translation of proteins, respectively [22]. These processes establish the diversity in the protein-coding potential of mRNAs. Specifically, RNA-binding proteins that regulate mRNA turnover and translation may have a substantial effect on age-dependent gene expression patterns [23]. A recent study has observed differing levels of expression of four turnover/translation RBPs in human fibroblasts during replicative senescence and tissue-level differences in expression from subjects varying in age [24]. One of these RBPs, AUF1, was detectable in the ovary. Another factor in the splicing of pre-mRNAs is the spliceosome, which is comprised of several small nuclear ribonucleoproteins [25]. During alternative splicing events, altered spliceosome expression seems to have an impact on oocyte maturation in human ovaries, allowing transcriptome profiles the opportunity for identifying biomarkers for oocyte quality [26]. Since maturation deficiencies are a significant symptom of ovarian aging, new methods that focus on the detection of the mechanisms responsible for the detrimental effects of aging need to be recognized [27].

This study aims to draw the attention of researchers in ovarian biology to the resolution of transcriptomic analysis. The basis is to view the transcript level of the transcriptome, analyze notable transcript variants, and determine whether their function is directly involved in the aging process. We observed significant transcripts that were differentially expressed in the aged ovary and those that were switched on and off in the aged ovary. Then, our discoveries have found transcripts in the primordial follicles of mature female mice that may impact the ovarian reserve, disrupting the regulation of both upstream and downstream processes. These observed transcripts have the potential to help find practices to stunt the ovarian aging process and prevent the harmful effects resulting from it.

## 2. Materials and Methods

### 2.1 Sample Collection and Processing

All procedures were performed in accordance with the protocols approved by the University of Kansas Medical Center Animal Care and Use Committee. At 6- and 12-months of age, age-matched wild-type (WT) female mice cytology was examined microscopically to determine reproductive cyclicity. On the day of the estrus cycle, the ovaries of the 6- and 12-month-old WT mice were collected immediately after euthanasia. One of the ovaries was then fixed in 4% paraformaldehyde (PFA), followed by histological examinations conducted on paraffin-embedded, hematoxylin and eosin-stained sections. The other ovary was snap frozen in liquid nitrogen and stored at -80°C until they were processed for total RNA analysis.

### 2.2. RNA Sequencing Data

This study includes RNA-Seq data from 6-month-old mouse ovaries (n=3 libraries) and 12-month-old mouse ovaries (n=3 libraries). Mouse ovary data were generated in the Stout laboratory at the Oklahoma Medical Research Foundation and have been downloaded from the Sequencing Read Archive (PRJNA1195555; SRA, NCBI).

### 2.3. RNA Sequencing Analysis

RNA-Seq data were analyzed using CLC Genomics Workbench 24 (Qiagen Bioinformatics, Redwood City, CA, USA). The software has Linux, Macintosh, and Windows versions; we used the Windows version to analyze the RNA-Seq data. CLC Genomics Workbench uses the expectation-maximization (EM) estimation algorithm to categorize and assign annotated transcripts to the transcript variants within the reference genome, gene, and mRNA. All clean reads were obtained by removing low-quality reads and trimming the adapter sequences. The high-quality reads were aligned to the Mus musculus reference genome (GRCm39), gene (GRCm39.113_Gene), and mRNA sequences (GRCm39.113_mRNA) using the default parameters: (a) maximum number of allowable mismatches was two; (b) minimum length and similarity fraction was set at 0.8, and (c) a minimum number of hits per read was 10. The expression values of individual gene identifiers (GE) or each mRNA splice variant (TE) in young and old ovaries were measured in TPM [24-26]. The threshold p-value was determined according to the false discovery rate (FDR). Differentially expressed genes were defined if the absolute fold change (FC) in expression was ≥ 2 with an FDR p-value of ≤0.05.

Of the 260,635 transcripts compiled in the reference genome, 5902 transcripts were differentially expressed in the old ovaries. We selectively analyzed 2779 TFs, curated by the Gifford lab from a list of human TFs, in addition to a list of 651 ERs from a gene atlas, 1296 RBPs from a comprehensive list of annotated RBPs, and 504 snRNPs from two articles covering spliceosomes [25,28-31]. Remarkably, about 90% of the transcripts observed in each group about aging ovaries expressed more than two transcript variants based on the GRCm39.113_mRNA reference build. New tracks containing only the specified groups were generated from each RNA-Seq data file containing GE or TE values, which were used in subsequent analyses. The threshold p-values were determined according to the FDR to identify the differentially expressed genes or transcript variants between young and old ovaries. A gene or a transcript variant was considered differentially expressed if the absolute FC was greater than or equal to two and the FDR p-value was less than or equal to 0.05.

### 2.4. Analysis of the Transcript Variants

We analyzed the differential expression of genes using the gene expression output of RNA sequencing analysis files. Here, all the mRNA splice variants are aggregated into one value. The differentially expressed genes were divided into three groups: upregulated (FC ≥2 and FDR p-value ≤0.05), downregulated (FC ≤ -2 and FDR p-value ≤0.05), and insignificant (either absolute FC <2 and/or FDR p-value >0.05). We analyzed the differential expression of transcript variants using the transcript expression output of the RNA sequencing analysis files. In contrast to the gene expression output, these files include each mRNA splice variant and its expression in the ovaries. The differentially expressed transcript variants were also divided into three groups: upregulated (FC ≥2 and FDR p-value ≤0.05), downregulated (FC ≤ -2 and FDR p-value ≤0.05), and insignificant (either absolute FC <2 and/or FDR p-value >0.05).

The nomenclature for the transcript variants follows the Ensembl transcript annotations, where each variant is called first with the gene name, followed by a hyphen, and then a three-digit number starting from 200. 201 and 202 are usually the Ensembl canonical, a single transcript chosen for a gene that is the most conserved, highly expressed, has the longest coding sequence, and has been discussed in many key resources like NCBI and UniProt. Nonetheless, the Ensembl canonical can vary from gene to gene, so it is important to reference the official Ensembl website for further details.

### 2.5. Statistical Analyses

For RNA Seq, each study group contained three library samples. In CLC Genomics Workbench, the ‘differential expression for RNA-Seq tool’ performs multi-factorial statistics on a set of expression tracks based on a negative binomial generalized linear model (GLM). The final GLM-fit and dispersion estimate calculates the total likelihood of the model given the data and the uncertainty of each fitted coefficient28. Two statistical tests-the Wald and the Likelihood Ratio tests-use one of these values. The Across groups (ANOVA-like) comparison uses the Likelihood Ratio test.

### 2.6. Gene/Transcript Switching Analysis

After the data had been analyzed, genes that showed no expression were extracted from the first sample of the control group. The resulting list of genes is then filtered in the following sample. After repeating the process for all samples, a list of genes that have no expression in all samples is the resultant output. The non-expressed list of genes was filtered in the experimental group to obtain a list of genes switched on due to ovarian aging. The exact process was done with the experimental and control groups to produce a list of genes switched off. The protocol for transcript switching matches that of gene switching.

## 3. Results

### 3.1. Changes in Aging Ovaries

The continual decrease in the number of primordial follicles and their quality poses a direct link to ovarian aging, and these processes weaken the ovarian reserve or cause premature ovarian insufficiency [32]. To understand the molecular mechanisms impacting follicle activity in the ovary, we underscored expressional changes in mRNA splice variants of various gene groups in young and old mouse ovaries. These expressional shifts gave insight into each transcript’s roles during ovarian aging. However, observing the phenotypic changes in an aging ovary precedes data analysis as it allows for the effects of senescence to come to view. Histology samples of young and aged ovaries were taken under a manual Nikon 80i brightfield microscope to record the follicular changes due to aging (**Figure 1**). After exploring the phenotypic development in the ovaries, we compared the RNA-Seq data of 6- and 12-month-old mouse ovaries and identified the differentially expressed transcript variants.

**Figure 1.**
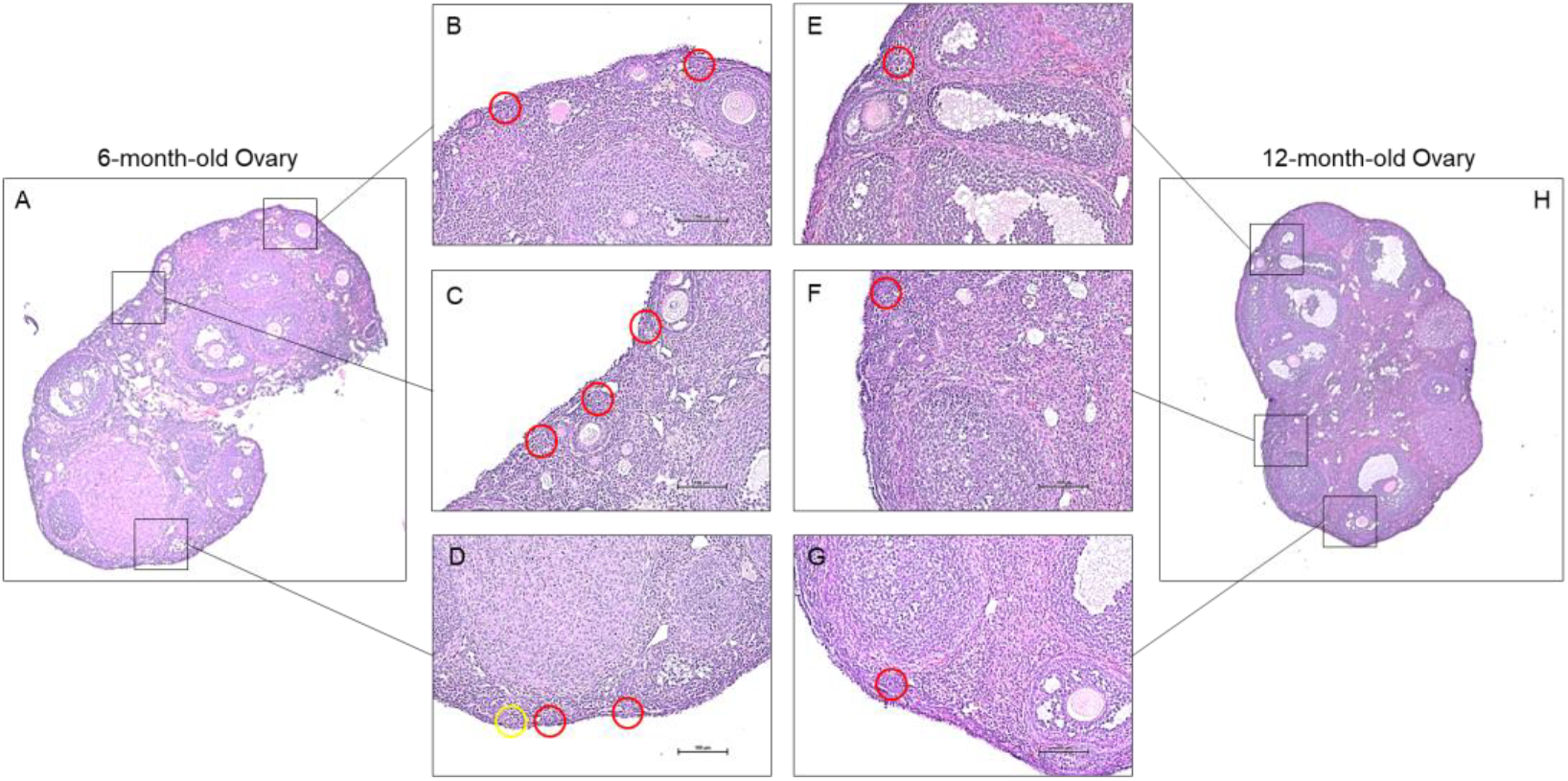
Histology of aging mouse ovaries. 6-month and 12-month control wild-type mice ovaries were isolated for histology sampling. The 6-month ovary (A) exhibited more primordial follicles than the 12-month ovaries (H). Sections were selected from the 6-month ovaries (B-D) to display multiple primordial follicles in the region of interest. The sections for the 12-month ovaries showed limited signs of primordial follicles and were difficult to identify (E-G). The red outlines indicate complete primordial follicles, and the yellow outline indicates a transitioning primordial follicle into a primary follicle.

### 3.2. Evaluation of RNA-seq Data

The RNA sequencing data was filtered through a FastQC quality control test before employing CLC Genomics Workbench’s and DNASTAR’s RNA sequencing analysis. Regarding per-base sequence quality, each fastq RNA sequencing file had a Phred score greater than 30, which was seen throughout all bases with no noticeable drop towards the ends of the reads. The per-sequence quality scores maintained a mean Phred score greater than 30, and the per-base sequence content displayed an even base composition for every position of the read. GC content was consistent throughout all sequences, per-base N content was very low, and sequence duplication levels were low. The only concern was in the overrepresented sequences, which indicated the presence of an adapter. These adapters were eliminated through a trimming process.

The RNA sequencing data were imported into CLC Genomics Workbench, where adapter cleaning and accurate sequencing were performed after trimming the reads within the software. RNA-seq results were cross-verified by both genomics programs and showed a promising arrangement along the first principal component axis. The two data sets, 6-month and 12-month ovaries, also displayed slight variation in their respective groups with similar averages, maximums, minimums, and outliers. Still, differences between both groups were seen in averages, maximums, and top outliers, where the 12-month transcripts were expressed at higher TPMs for gene and transcript expression (**Figure 2A-D**). However, due to the number of genes compared to transcripts, the transcript expression plots had tighter averages, balanced by a larger number of outliers. The quantities between genes and transcripts were similar when viewing TPM and FC in a vacuum (**Figure 2E-H**).

**Figure 2.**
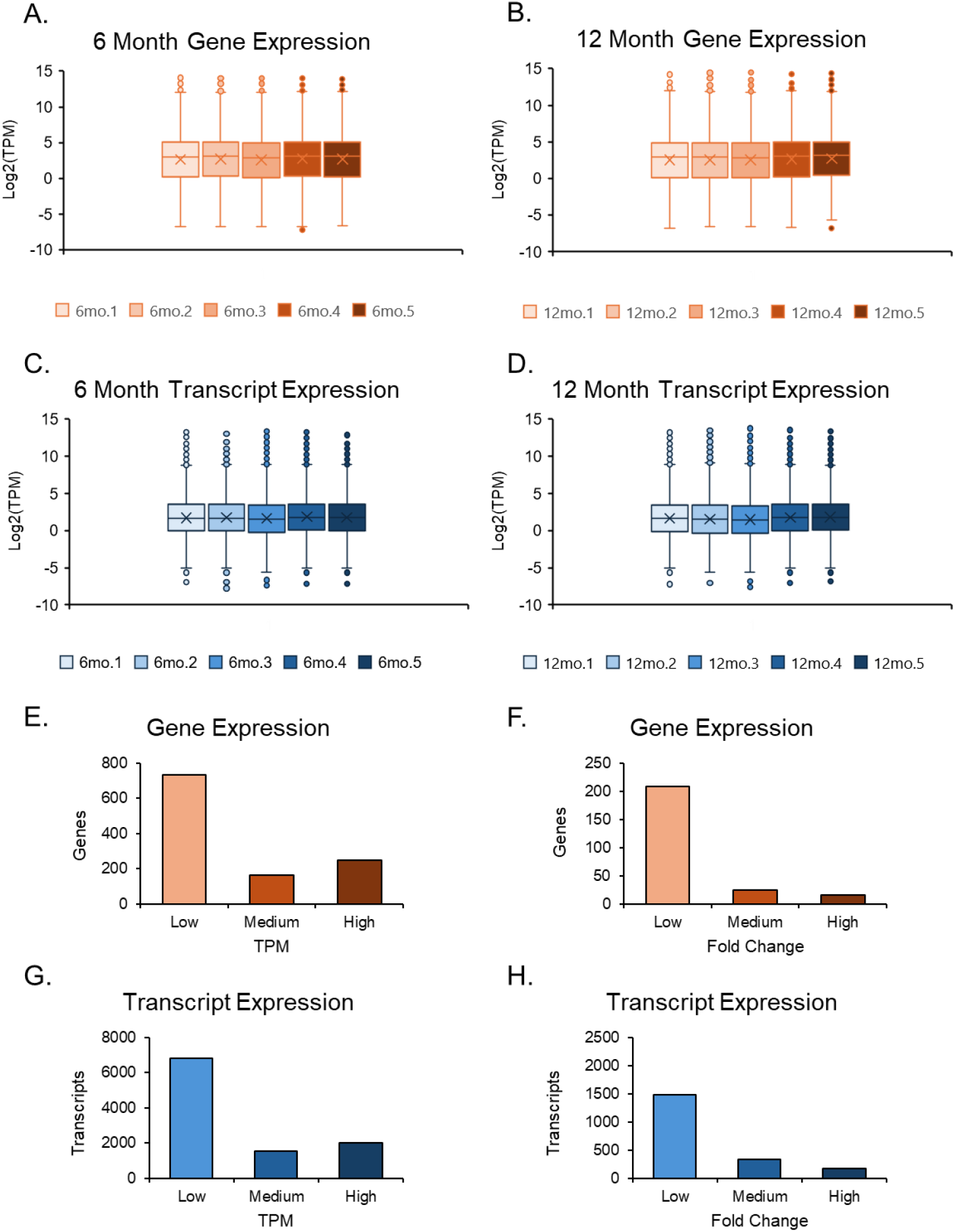
Expression plots of the overall data set. Box and whisker plots of both 6-month and 12-month data sets were separated; their gene and transcript expression values were evaluated after normalization. The 6-month expression data (A, C) among 5 samples were compared. The 12-month expression data (B, D) were compared similarly to the 6-month data. Gene and transcript expression was assessed further by separating TPM values into three groups: low (1< TPM ≤5), medium (5< TPM ≤10), and high (TPM >10) (E, G). Additionally, the same form of analysis was done for FC values by separating them into three groups: low (2< FC ≤5), medium (5< FC ≤25), and high (FC >25) (F, H).

### 3.3. Differential Expression of Genes in Aging Ovaries

After the RNA sequencing analysis, a gene expression output was observed in the differential expression of genes in aging ovaries. Compared to the ovaries from 6-month-old young mice, the ovaries from 12-month-old middle-aged mice showed 337 differentially expressed transcript variants out of 64,470 genes compiled in the GRCm39 reference genome (**Figure 3A**). Of those, 182 genes were upregulated, and 155 were downregulated (absolute FC ≥2 and FDR p-value ≤0.05) with a TPM larger than 5.

**Figure 3.**
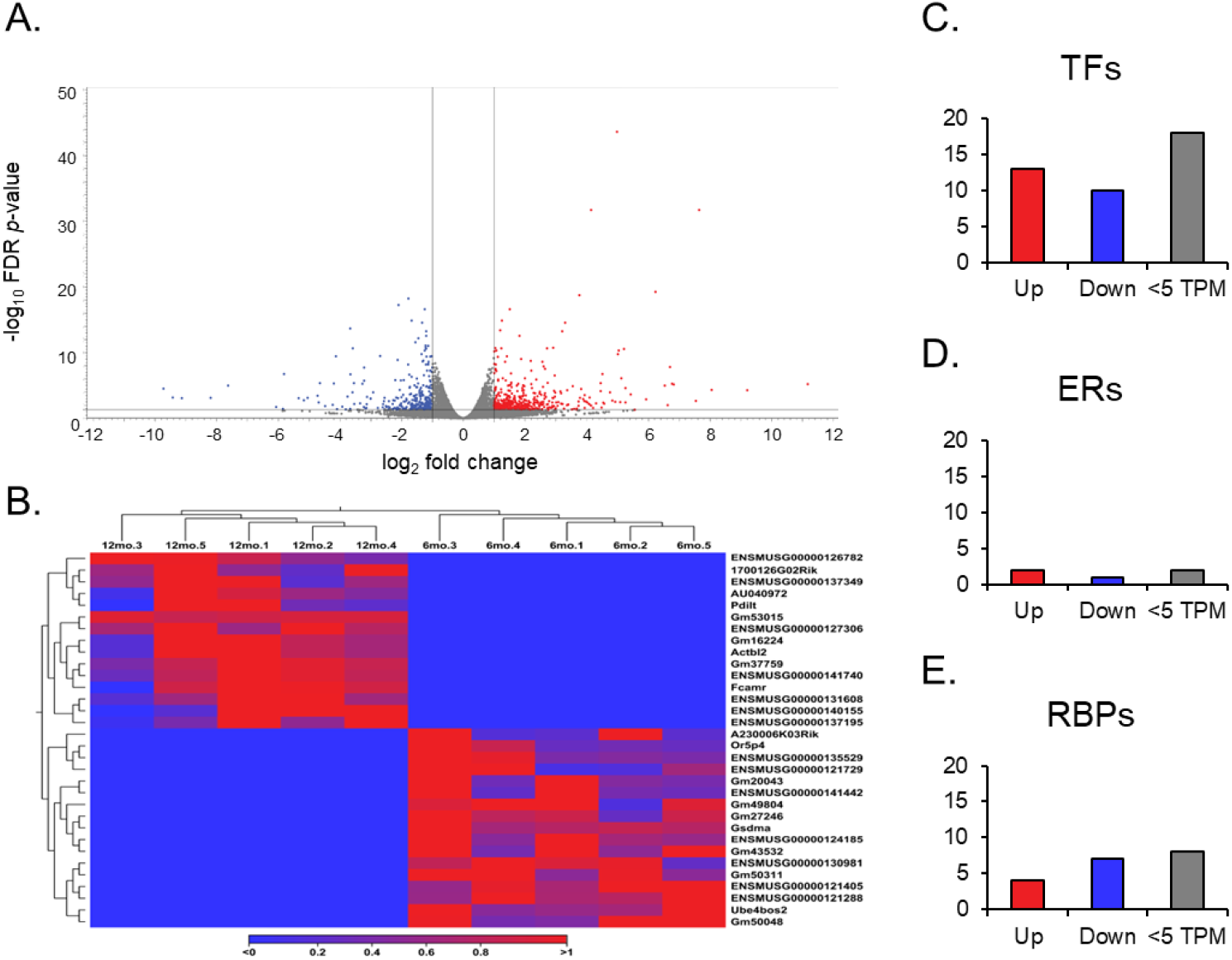
Gene expression volcano plot, gene switching heat map, gene group analysis. The volcano plot displays all expressed genes (A) when comparing the middle-aged and young ovaries. The genes with the low FC and high p-value points faded out. The red and blue points represent differentially expressed genes, red points indicate upregulated and blue indicate downregulated genes in middle-aged ovaries compared to young ovaries. Heat map for the expression profiles of all genes switched off and on (B) either in young or middle-aged mice ovaries. The expression levels were visualized with a gradient color scheme where the right side of the bar, red, represents high expression, and the left side, blue, represents low expression relative to the data set. Significant genes from each group, transcription factors (TFs), epigenetic regulators (ERs), and RNA binding proteins (RBPs) were sorted into upregulated (FC ≥2), downregulated (FC ≤2), and low TPM (<5) groups (C-E).

Further analysis of expression results allowed for the observation of gene switching (**Figure 3B**). Genes that exhibited no expression across all data entries in one group, either at 6-months or 12-months, were filtered in the opposite group for expression to identify genes that have been switched on (no expression in the young ovaries, expression in the aged) and off (no expression in the aged ovaries, expression in the young ovaries) during the aging process in the ovary. Of the 17 genes switched off, only one, *Gm49804*, was highly expressed (TPM >10) in the young ovary. Of the 15 genes switched on after aging, *Gm53015* was the only highly expressed (TPM >10) in the aged ovary. In addition, we analyzed differentially expressed genes with a TPM greater than 5, and found 23 transcription factors (TFs), 3 epigenetic regulators (ERs), and 11 RNA-binding proteins (RBPs) (**Figure 3C-E**).

No snRNPs from the curated list were significant enough to be detected. Out of the 23 TFs, 13 were significantly upregulated, specifically, *Zbed6* and *Sfrp4* were noted due to their high FC (>30) and expression value (TPM >2000), respectively. Of the 10 downregulated TFs, *Nr1d1, Osr2, Dbp*, and *Hspa1b* exhibited TPMs above 100. However, in the ERs, only two genes were significantly upregulated, and one gene was significantly downregulated. Notably, *Phf20l1*, which is upregulated, and *Spen*, which is downregulated, were distinguishable since they had TPMs above 30 in that small group. For RBPs, we detected 4 significantly upregulated genes. However, none exhibited any marked characteristics. On the other hand, 7 genes were significantly downregulated, where the aforementioned, notable genes, *Spen* and *Gm49804*, were detected with *Nynrin*, a gene with a TPM greater than 30.

### 3.4. Differential Expression of Transcript Variants in Aging Ovaries

A transcript expression output follows a gene expression output during an RNA sequence analysis. The transcript expression produced the necessary information to understand the differential expression of splice variants in aging ovaries. When looking at each mRNA splice variant within the GRCm39 reference genome, the expression comparison between young and aged ovaries became more in-depth as the genome compiles 260,635 total transcripts (**Figure 4A**). Of the vast amount of transcripts, 2,510 were significant during differential expression. 1,264 of the significant transcripts were upregulated (FC ≥2 and FDR p-value ≤0.05), and 1,241 were downregulated (FC ≤ -2 and FDR p-value ≤0.05), both with a TPM larger than 5.

**Figure 4.**
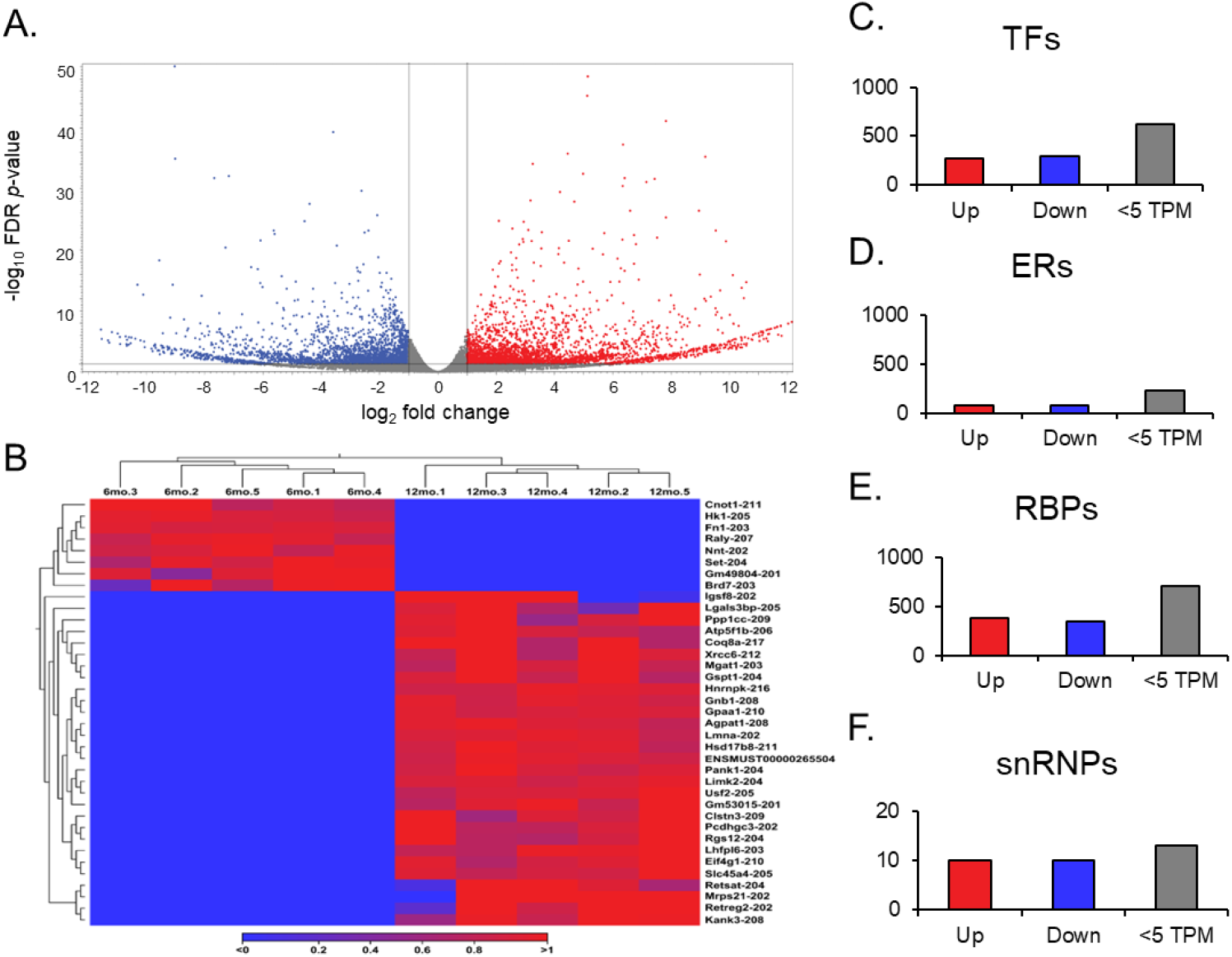
Transcript expression volcano plot, transcript switching heat map, transcript group analysis. The volcano plot displays all expressed transcripts (A) when comparing the middle-aged and young ovaries. The transcripts with the low FC and high p-value points faded out. The red and blue points represent differentially expressed genes, red points indicate upregulated and blue indicate downregulated genes in middle-aged ovaries compared to young ovaries. Heat map for the expression profiles of all transcript variants switched off and on (B) either in young or middle-aged mice ovaries. The expression levels were visualized with a gradient color scheme where the right side color, red, was used for high expression, and the left side color, blue, was used for low expression relative to the data set. Significant transcripts from each group, transcription factors (TFs), epigenetic regulators (ERs), RNA binding proteins (RBPs), and spliceosomes (snRNPs) were sorted into upregulated (FC ≥2), downregulated (FC ≤2), and low TPM (<5) groups (C-F).

Gene switching was observed for additional analysis, but at the transcript level (**Figure 4B**). The transcript variants that showed no expression across all data entries in one group, 6 months or 12 months, were filtered in the opposite group for expression to identify genes switched on and off during the aging process in the ovary. Out of the 105 transcript variants switched off after aging, 8 were highly expressed in the young ovary. Notably, *Raly-207* and *Set-204* had the most significant difference, having TPMs greater than 20 before getting switched off in the aged ovary. Of the 158 transcript variants switched on after aging, 30 were highly expressed in the aged ovary. Here, *Eif4g-210, Gnb1-208, Hsd17b8-211*, and *Ppp1cc-209* displayed TPMs greater than 30 when switched on in the aged ovary, and *Atp5f1b-206* was expressed at a TPM above 150.

We observed different groups of differentially expressed transcripts with TPM values above 5 (**Figure 4C-F**). We have detected 569 TFs, 149 ERs, 728 RBPs, and 10 snRNPs. Among the TFs, 272 significantly upregulated transcripts were found. *Sfrp4-204, Tsc22d1*-*214*, and *Nfe2l1-212* were notable transcripts within the upregulated group due to their simultaneous features of a high TPM (>100) and high FC (>10). Although the other 297 significantly downregulated group of transcripts did not exhibit the same magnitude of expression as their upregulated counterparts, *Raly-207* and *Set-204* from the transcript switching analysis reappeared as notable transcripts due to their TPM and FC compared to other downregulated transcripts. For the group of ERs, 72 were significantly upregulated, and 77 were significantly downregulated. Within the upregulated group, *Ppm1g-205* and *Dek-204* stood out due to their high FC (>1000), but *Ncl-205* and *Ogt-203* also stood out due to their high TPM (>40). *Usp3-211* remained in the downregulated group with an FC above 1000, and Ywhaz-204 had an expression value above 100 TPM. To analyze the RBPs, we followed a similar guideline as the TFs due to their comparable transcript quantities. 379 RBP transcripts were detected as significantly upregulated, and 349 were detected as significantly downregulated. *Actg1-204, Atp5f1b-206, Rplp2-203, Rack1-202, Psap-202, Eef1g-205* are noted for their high TPM (>100) and high FC (>10) in the upregulated group. *Rps3-205* and *Rpl13a-214*, on the other hand, are regarded for the same quantitative measures but in the downregulated group. Finally, the snRNPs only contained 10 significantly upregulated transcripts and 10 significantly downregulated transcripts. Of those 10 upregulated transcripts, *Slu7-206* stood out for the large FC (>1000), but *Sf3b2-208* and *Snrnp70-209* are noted for their high expression values (TPM >50). The 10 downregulated transcripts do not have impressive quantitative results, but *Sf1-203* was the most downregulated of the group. It was notable for being a transcript of *SF1*, an essential gene for the development and function of the ovary.

### 3.5. Correlation of Differentially Expressed Genes and Transcript Variants

Although the same genes and their corresponding transcripts were present in the GRCm39 reference genome, further analysis demonstrated a notable discrepancy between gene expression and transcript expression analysis. A record of each splice variant of the original 337 significant genes was kept to analyze such discordances. This resulted in 1,285 transcript variants, where only 241 were statistically significant. For confirmation, these 241 significant variants were the only variants to match the original 2,505 significant variants from the transcript expression output. Of these 241 significant variants extracted from gene expression, 125 were upregulated (FC ≥2 and FDR p-value ≤0.05), and 116 were downregulated (FC ≤ *-2* and FDR p-value ≤0.05), both with a TPM value larger than 5.

Discrepancies in gene and transcript switching follow the logic of varying expression of multiple splice variants within one gene. By taking the genes that have switched on and off in the aging ovary and selecting their respective transcripts, a heat map was created to display the genes involved in gene switching at the transcript level (**Figure 5B**). This process was done for the transcripts involved in transcript switching by taking the transcripts and making a heat map of their respective genes to show transcript switching at the gene level (**Figure 5A**).

**Figure 5.**
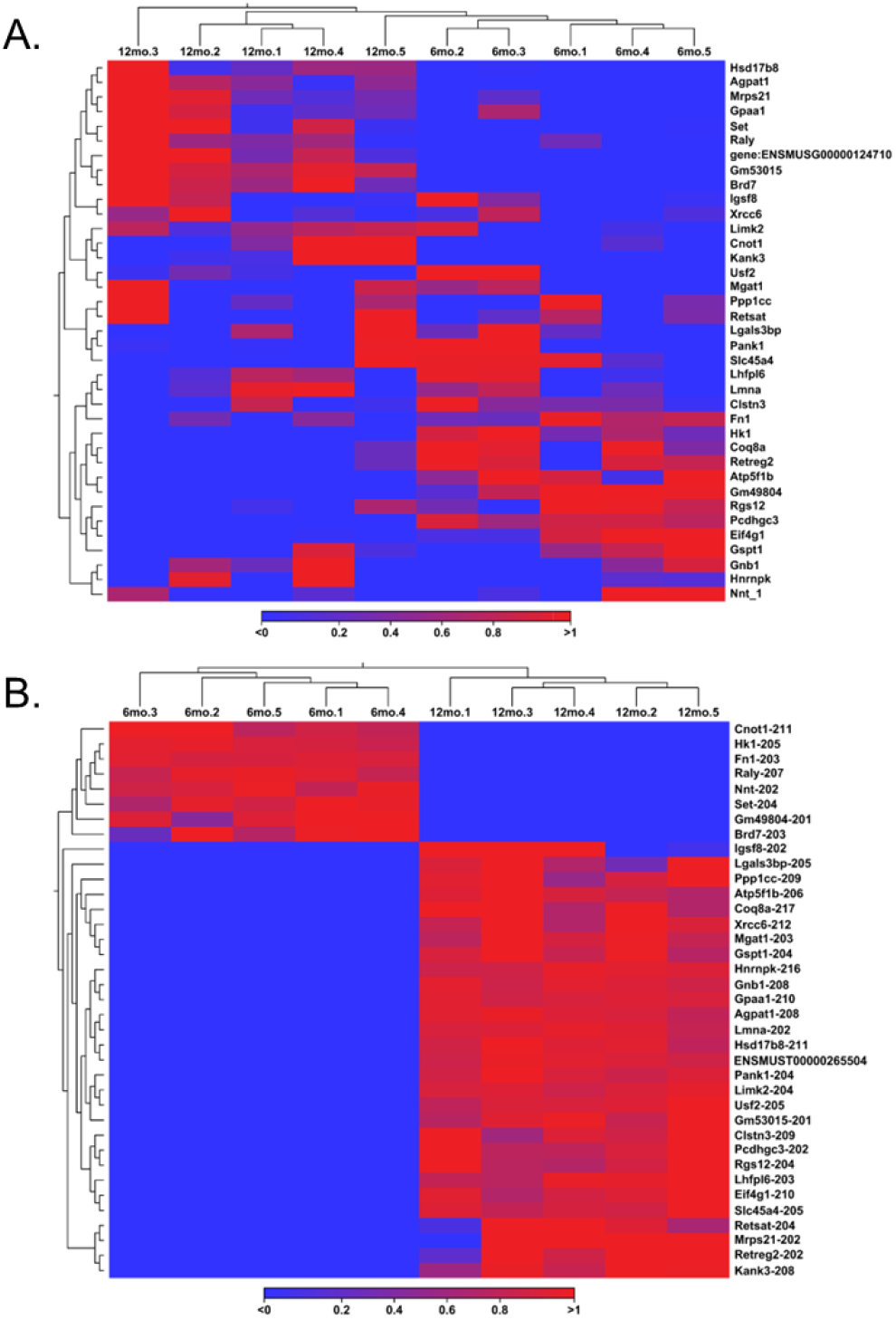
Switching discrepancies heat map. Transcripts that participated in transcript switching were filtered into a gene expression analysis to isolate each gene from which the transcripts originated (A). Genes that participated in gene switching were taken and filtered into a transcript expression analysis to pull each transcript related to those genes (B).

### 3.7. Mechanisms Underlying Altered Transcript Variants in Aging Ovaries

After analyzing each group individually, we examined the relationships between related groups, such as TFs compared with ERs and RBPs compared with snRNPs. Overlaps were detected in the mentioned comparisons, which will help identify transcripts that play multiple essential roles. When comparing TFs to ERs, there were many overlaps between the two groups, as many transcripts share an upstream role in transcription and epigenetics (Figure 6A). Of these overlapped transcripts, four were remarkable due to their relation to the ovary and high expression value when differentially expressed. For instance, the transcript Npm1-203 had a high TPM value (TPM >50) and a remarkably downregulated FC (<-20). An overexpression of NPM1 is linked to unsuitable outcomes for women with ovarian cancer [33]. Mta1-202 was also a transcript that was highly down-regulated (FC <-11) and expressed at a high level (TPM >40). MTA1 is associated with advanced and metastatic ovarian cancer tissue [34]. In contrast, the transcript, Trim28-203, originated from a gene recently found to regulate granulocyte senescence [35]. Here, our analysis indicated that the transcript was expressed at a high TPM (>40) and upregulated heavily (FC >10). The final transcript, Ube2n-203, was also upregulated to a high degree (FC >5) and expressed highly (TPM >40). The gene UBE2N is seen to be key in the regulation of paclitaxel, a chemotherapeutic drug, resistance in ovarian cancer cells [36].

**Figure 6.**
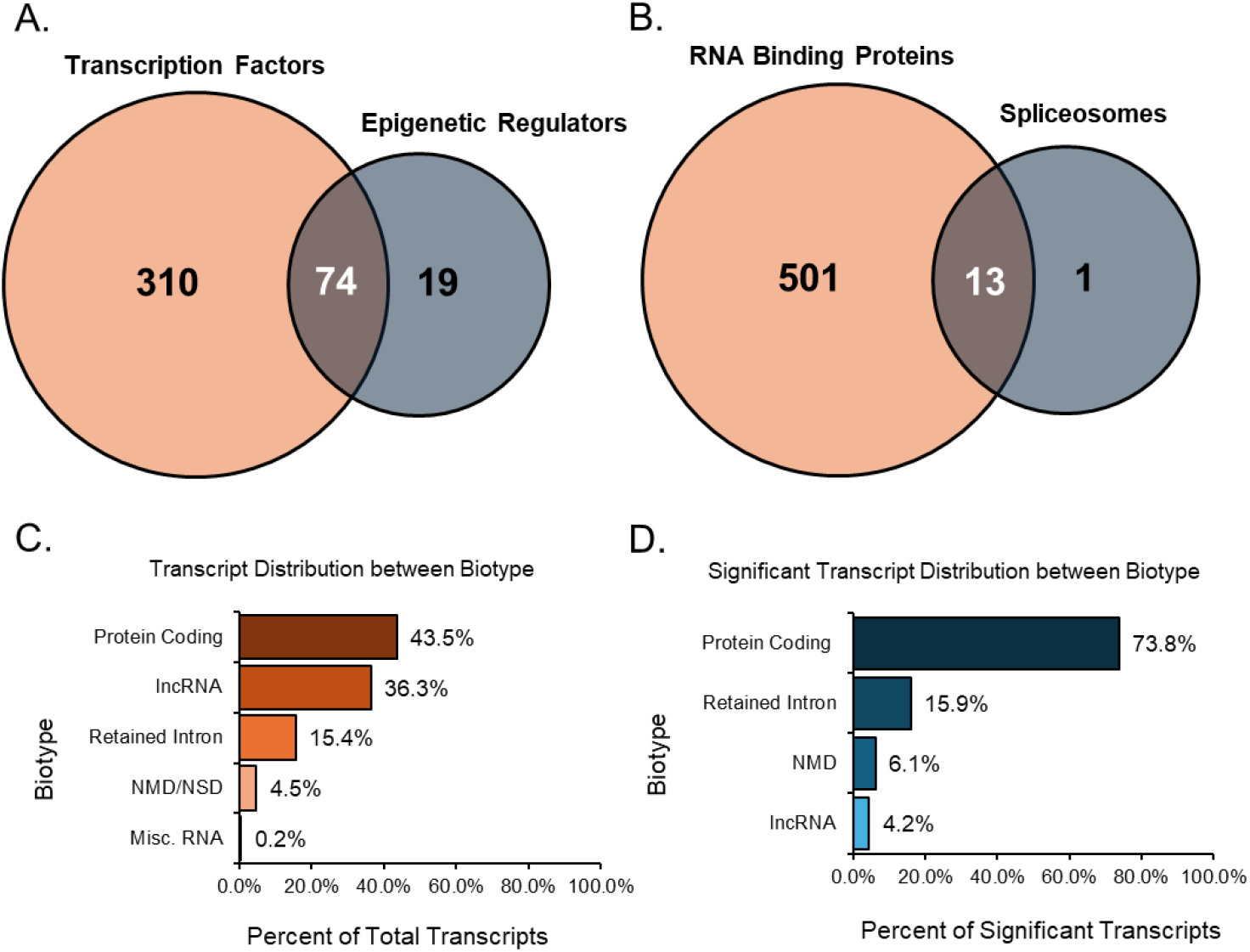
Venn Diagram of Transcript Groups and Transcript Variant Biotype Distribution Graph. The first Venn diagram quantified significant (TPM >10, absolute FC ≥2, and FDR p-value ≤0.05) transcript factors and epigenetic regulators (A) and found the transcripts that both groups share. Similarly, the second Venn diagram quantified significant RNA binding proteins and transcripts associated with spliceosomes (B) while finding the transcripts in both groups. All expressed transcript variants from the data were categorized by their biotype (C), giving a percentage distribution of all groups. Significant transcripts from the data were categorized by their biotype (D), giving a percentage distribution of their resultant groups.

Significant transcripts associated with RNA binding proteins and spliceosomes were compared, and we found 13 unique transcripts that overlapped between these two groups (**Figure 6B**). Of these 13 transcripts, *Rbm25*, known as a global splicing factor, appeared as two splice variants (-206, -208). Both variants were upregulated and displayed high TPM values over 10 and FC values over 5. Similarly, *Sf3b1-203*, a disruptive splicing factor, was upregulated in both groups with high TPM and FC values. In contrast, *Cd2bp2-201* was a variant formed via alternative start sites and was highly downregulated in both groups. Despite not being the full-length canonical variant of the *CD2BP2* line, *Cd2bp2-201* still coded for the same protein and was the only significant variant present during analysis.

Most of the expressed, annotated transcript variants detected in the study were of the protein-coding biotype. These protein-coding transcript variants contain an open reading frame that can be translated into a protein, and they make up 35.5% (45,561 transcripts) of all expressed transcripts (**Figure 6C**). In addition, 7.9% (10,229 transcripts) of all expressed transcripts were alternative splice variants of protein-coding transcripts without a defined coding sequence. The protein-coding transcripts comprised 43.5% (55,790 transcripts) of all transcripts. Following the protein-coding biotype, long non-coding RNA (lncRNA) comprised 36.3% (46,548 transcripts) of all expressed transcripts. These lncRNAs did not contain meaningful open-reading frames but were longer than 200 base pairs. Another notable biotype included those involved in decay processes such as non-sense-mediated decay (NMD) and non-stop decay (NSD). The transcripts classified as nonsense-mediated decay had a premature stop codon, most likely subjected to targeted degradation. For Ensembl, nonsense-mediated decay prediction happened when an inframe termination codon was found more than 50 base pairs upstream from the final splice junction. Transcripts related to decay made up 4.5% (5,809 transcripts) of the expressed transcripts in the study. Intron retention was another alternative splicing outcome, where the intron was kept rather than spliced out. Transcripts categorized as retained introns were splice variants with similar intronic sequences to other transcripts produced from the same gene locus; these transcripts comprised around 15.4% (19,807 transcripts) of all expressed transcripts. Finally, miscellaneous RNA will be a combination of non-coding RNAs unable to be classified, RNA components of a ribosome, snRNAs, snoRNAs, miRNAs, and mitochondrial RNAs, making up 0.2% (266 transcripts) of all expressed transcripts.

After filtering the significant transcripts from the expression data with a minimum TPM threshold of 5, an absolute FC of at least 2, and an FDR p-value of 0.05 or lower, we identified the biotype of all resultant transcripts (**Figure 6D**). Here, protein-coding transcripts, including those without a defined coding sequence, comprised 73.8% (1853 transcripts) of significant transcripts. Interestingly, the next largest group of biotypes was the retained intron transcripts, which comprised 15.9% (400 transcripts) of the differentially expressed transcripts. Non-stop decay transcripts were not detected as significant, so only NMD was recorded. These decay transcripts comprised 6.1% (152 transcripts) of substantial transcripts. Despite being the second largest biotype in the total transcript analysis, lncRNA made up 4.2% (105 transcripts) of significant transcripts. The shift from lncRNA to more protein-coding transcripts may give insight into the mechanisms involved in the ovarian aging process.

## 4. Discussion

Nearly half of the global population experiences a range of endocrine, metabolic, and neurological disorders as they enter the later stages of life, with a significant portion of these issues stemming from ovarian aging [37]. The ovaries are among the first organs to show functional decline, particularly in reproductive capacity [38]. This has profound implications for women’s health and well-being. As the ovaries age, they undergo a series of complex changes that impact hormonal balance, tissue integrity, cellular function, and gene regulation [4]. Despite recently developed therapies aimed at mitigating the symptoms of ovarian aging, such as hormone replacement therapies (HRT), these treatments often come with significant side effects, raising concerns about their safety and efficacy for long-term use [39]. A comprehensive understanding of the cellular and molecular mechanisms underpinning ovarian aging is essential for devising targeted interventions to enhance women’s health outcomes.

Various molecular hallmarks of aging contribute to the decline in ovarian function [40]. These hallmarks include genomic instability, which refers to the increased likelihood of mutations; telomere attrition, which is the progressive shortening of protective chromosome ends; epigenetic alterations that affect gene expression without changing the DNA sequence; loss of proteostasis, which disrupts the balance of protein synthesis and degradation; dysregulation of nutrient sensing pathways, mitochondrial dysfunction that affects energy production; cellular senescence, which is the process by which cells lose their ability to divide; stem cell exhaustion, leading to a reduced capacity for tissue regeneration; and altered intercellular communication, which can disrupt the signaling pathways necessary for maintaining ovarian health [41-49]. Using bulk RNA-sequencing, differentially expressed genes in aging mammalian ovaries can be identified. We have conducted an in-depth analysis of gene regulation at the transcript level, providing insight into the actual changes in the ovarian transcriptome. Our analysis has uncovered significant differences in the expression of various transcript variants within the ovary; this phenomenon is not exclusive to rodents, but likely extends to humans and primates as well. Furthermore, the expression of these transcript variants is linked to upstream regulatory mechanisms involving epigenetic and transcription factors. Meanwhile, downstream processes, such as RNA modifications mediated by RNA binding proteins and spliceosomes play a crucial role in shaping the final output of gene expression.

Observing significant differences in aging ovaries can be challenging, as our expression plots revealed minimal variation across different age groups. However, histological analysis revealed a nuanced picture, demonstrating notable phenotypic changes. Specifically, young ovaries are typically characterized by a substantially higher number of primordial follicles than middle-aged mice. This discrepancy in follicular quantity is likely a result of the transcriptomic changes that occur as ovaries age, which can alter the molecular landscape of ovarian function and development. A more thorough examination of the transcripts involved can yield a more accurate depiction of the aging process, due to the enhanced quantity and resolution that modern transcriptomic analyses provide. Given the focus of the current study on quantitatively assessing specific gene groups about aging, the choice of software for data analysis is paramount. CLC’s genomics software stands out for its use of a specific aligner algorithm, the expectation-maximum algorithm, which has been demonstrated to perform consistently with high accuracy. When benchmarked against other popular aligners, such as STAR and NOVOALIGN, CLC has been recognized for its performance, especially when analyzing data at the “junction level” [50]. This level of precision is crucial for understanding the complexities associated with gene expression in aging ovaries. Consequently, this study predominantly employs CLC, as it demonstrates top performance across all data sets, except for those that were the least complex, where complexity is determined by the difficulty of alignment in a specific region.

Gene expression analysis was initially done to extract as much information as possible, despite the challenge of aggregating all transcripts into a single gene identifier. Out of the 64,470 genes in the reference genome, 38,159 were expressed. However, using a stringent filtering process, we discovered that only 337 met our established criteria denoting statistical significance. These criteria were a minimum TPM threshold of 5, an absolute FC of at least 2, and an FDR p-value of 0.05 or lower. Among the different groups analyzed, we found that transcription factors (TFs) yielded 24 significant genes, approximately 7% of the total significant genes identified in this study. In stark contrast, epigenetic regulators (ERs) yielded a mere 3 significant genes, representing only 0.9% of the significant gene pool. It is important to note that the overlap between TFs and ERs yielded a combined 24 distinct genes, highlighting the interconnected nature of these regulatory mechanisms. This suggests that approximately 7% of the genes highlighted in this study play some role in transcriptional regulation or epigenetic modifications. The group comprising RNA-binding proteins (RBPs) and spliceosome components (snRNPs) revealed only a modest enhancement over the contributions of TFs and ERs. A total of only 11 significant genes were identified from the RBPs, while snRNPs did not contribute any significant genes to the analysis. This group accounted for approximately 3% of the significant genes identified in this study. The gene switching analysis, aimed at identifying genes with completely altered expression patterns, identified a limited number of genes. Among these, only two genes exhibited a TPM greater than 10: *Gm53015*, which was found to be switched on in the aged ovary, and *Gm49804*, which was switched off. Despite the seemingly low involvement of each group in the overall gene expression analysis, several notable genes emerged due to their remarkably high expression levels and significant FCs when comparing values from young to aged ovaries. Specifically, the genes *Zbed6, Sfrp4*, and *Phf20l1* demonstrated substantial upregulation, indicating their potential importance in the aging process in ovarian tissues. Conversely, genes such as *Nr1d1, Osr2, Dbp, Hspa1b*, and *Spen* showcased significant FCs and elevated TPM values despite being downregulated. These findings underscore the complexity of gene expression dynamics and highlight key players that may be pivotal in understanding the molecular underpinnings of ovarian aging.

A more detailed transcript expression analysis was conducted following the gene expression analysis, yielding more promising outcomes. Out of the 260,635 transcripts available in the reference genome, 128,220 were expressed, resulting in the detection of 2,510 significant transcripts (TPM ≥5, absolute FC ≥2, and FDR p-value ≤0.05). At the transcript level, 569 significant transcripts were identified, representing approximately 23% of the significant transcripts detected in this study. Notably, the number of significant ERs increased to approximately 6% for transcripts with 149 detected. Following the removal of overlapping transcripts and the subsequent compilation of a list of TFs and ERs, 604 significant transcripts were identified. This indicated that approximately 24% of the significant transcripts found in this study were associated with TFs or ERs, indicating their potential in the regulation of transcriptional processes or epigenetic modifications, an impressive enhancement compared to the initial data derived solely from gene expression analyses. For RBPs, a total of 728 significant transcripts were identified, which accounted for 29% of the overall significant transcripts in the study. The snRNPs also experienced a notable increase, with 20 significant transcripts detected. Although snRNPs only accounted for approximately 0.8% of the significant transcripts, the opportunity to investigate these 20 specific transcripts in relation to their roles in ovarian aging is considerably more feasible than conducting a broad gene-level analysis, particularly since no snRNPs emerged as significant at the gene level. When compiled, RBPs and snRNPs together still constitute approximately 29% of the significant transcripts identified in this study. This suggests that 731 transcripts may possess the capacity to influence downstream biological processes through the modifications they impose on RNA, especially in the context of ovarian aging. Switching analysis proved to be more effective than the gene switching analysis, revealing 30 transcripts that were activated in the aged ovary, each exhibiting a TPM greater than 10, alongside 8 transcripts that were switched off. Each of these highly expressed transcripts exhibited overlap with at least one of the four groups used in the expression analysis. Specific transcripts such as *Atp5f1b-206, Eif4g1-210, Gnb1-208, Hsd17b8-211*, and *Ppp1cc-209* are recognized for their high expression when switched on, while *Raly-207* and *Set-204* are highlighted for their high expression despite downregulation in the aged ovary. This nuanced understanding of transcript dynamics in the context of ovarian aging presents a valuable opportunity for further exploration.

Despite the vast number of long non-coding RNAs present within the dataset, a significant majority of the identified transcripts were classified as protein-coding. These transcripts harbor open reading frames (ORFs) that allow for protein translation, ultimately leading to the manifestation of specific biological functions [51]. Although many transcript variants code for the same protein, some may diverge and encode proteins with differences in sequence, differences in length, inclusion or exclusion of various domains, and even potential differences in biological function (e.g., enzymatic activity, binding capability, etc.) [49]. This variability in coding sequences among transcripts emphasizes the critical need to examine individual transcripts and their unique contributions to the transcriptome. Generalizing a transcript to its gene identifier may mask its true functional significance [52]. Accordingly, gene and transcript expression analysis often yield divergent outcomes, reflecting the inherent differences in the methodologies employed. Gene expression analysis provides a broader and more rapid overview of the dataset, effectively summarizing the expression levels of the aggregated transcripts. Conversely, the more detailed approach of transcript expression analysis permits a more thorough evaluation of individual transcripts, allowing a deeper understanding of expression patterns. By looking at transcripts in isolation, more expression data can be pulled, and more accurate depictions of gene expression values can be drawn. It is crucial to recognize that the expression of a single transcript cannot be reliably used as a proxy for the expression of a gene, nor can the reverse be assumed. Either gene expression or transcript expression analysis has its advantages and disadvantages. However, this study prioritizes the value of transcript expression in understanding the complexities of aging ovaries.

Throughout the study, transcripts from the genes *Gnb1, Sf1*, and *Sfrp4* were recognized for their dramatic shifts in expression in the aged ovary. A recent study showed *GNB1’s* potential involvement in polycystic ovary syndrome (PCOS) through single-nucleotide polymorphism analysis [53]. Our analysis demonstrates that *Gnb1’s* transcript, *Gnb1-208*, was switched on in the aged ovary at a high expression value (TPM >10). According to the Ensembl database, *Gnb1-208* codes for the same protein as its canonical counterpart, sharing functional relevance, but was expressed differently since *Gnb1-203*, the canonical transcript, was downregulated. Altering the expression of *Gnb1-208* may delay or prevent PCOS, ultimately halting a symptom of an aging ovary.

*S*teroidogenic factor 1 (*SF1*), has been demonstrated to be an important element in the establishment and maintenance of the ovarian reserve [54]. Our analysis identified the splice variant, Sf1-203, as the most significantly downregulated transcript within the spliceosome group. Notably, *Sf1-203* does not encode the same protein as its canonical counterpart, *Sf1-208*, indicating that these two variants fulfill distinct biological roles. *Sf1-203* is essential for the splicing of precursor mRNA, whereas *Sf1-208* is necessary for the ATP-dependent initiation of spliceosome assembly [51]. The stability of the canonical transcript’s expression suggests that manipulating the splice variant expression, *Sf1-203*, could be a strategic approach for preserving the ovarian reserve.

Finally, *SFRP4* has been characterized as a negative regulator of ovarian follicle development, limiting the ovarian reserve and reducing female fertility [55]. Like the *SF1* transcripts, the variant *Sfrp4-204* was highly upregulated, which does not produce the same protein as its canonical counterpart, *Sfrp4-203*. However, both transcripts were up-regulated in our analysis, and lower expression of *Sfrp4* can increase the ovarian reserve for female fertility. These transcripts associated with ovarian follicle dynamics may serve as critical molecular indicators that could delay the detrimental effects of ovarian aging through targeted expression modifications. Targeting the functional decline of the aging ovary may represent an optimal strategy for mitigating the impact of ovarian senescence towards the stagnation of aging ovaries, and the modulation of transcript expression could provide a foundational approach for advancing this field of research.

## 5. Conclusions

We analyzed the transcriptomes of young and aged mouse ovaries to identify potential gene changes during ovarian aging. Instead of the canonical ‘one gene one mRNA’ strategy, we have identified the changes in transcript variants in aging ovaries. Our analyses identified remarkable changes in transcript variants during ovarian aging in mice. To understand the mechanisms underlying altered transcript variants, we focused on key regulatory genes, including transcription factors, epigenetic regulators, RNA-binding proteins, and spliceosome components. The implications of our findings on transcript variation are profound, as they may directly influence the quantity and quality of oogenesis and steroidogenesis, thereby affecting female fertility and women’s health. Moreover, identifying these transcripts and their presumed biological functions provides valuable insights into the molecular underpinnings of ovarian aging.

## Author Contributions

M.R. conceptualization, supervision, and resources; K.V. data curation, methodology, investigation, formal analysis, and original draft preparation; G.P., R.T., and P.E.F. reviewing and editing, making a substantial contribution in preparation of the manuscript. All authors have read and agreed on the contents of the manuscript.

## Data Availability Statement

SRA, NCBI.

## Acknowledgements

We acknowledge the editorial board of Cells for waiving the publication fees and Qiagen Bioinformatics for their continued support.

## Disclaimer/Publisher’s Note

The statements, opinions and data contained in all publications are solely those of the individual author(s) and contributor(s) and not of MDPI and/or the editor(s). MDPI and/or the editor(s) disclaim responsibility for any injury to people or property resulting from any ideas, methods, instructions or products referred to in the content.

## References

1. Wu, J.; Liu, Y.; Song, Y.; Wang, L.; Ai, J.; Li, K. Aging conundrum: A perspective for ovarian aging. Front Endocrinol (Lausanne) 2022, 13, 952471, doi:10.3389/fendo.2022.952471.

2. Santoro, N.; Banwell, T.; Tortoriello, D.; Lieman, H.; Adel, T.; Skurnick, J. Effects of aging and gonadal failure on the hypothalamic-pituitary axis in women. Am J Obstet Gynecol 1998, 178, 732–741, doi:10.1016/s0002-9378(98)70483-1.

3. Weiss, G.; Skurnick, J.H.; Goldsmith, L.T.; Santoro, N.F.; Park, S.J. Menopause and hypothalamic-pituitary sensitivity to estrogen. Jama 2004, 292, 2991–2996, doi:10.1001/jama.292.24.2991.

4. Yan, F.; Zhao, Q.; Li, Y.; Zheng, Z.; Kong, X.; Shu, C.; Liu, Y.; Shi, Y. The role of oxidative stress in ovarian aging: a review. Journal of ovarian research 2022, 15, 100.

5. Revenkova, E.; Herrmann, K.; Adelfalk, C.; Jessberger, R. Oocyte cohesin expression restricted to predictyate stages provides full fertility and prevents aneuploidy. Curr Biol 2010, 20, 1529–1533, doi:10.1016/j.cub.2010.08.024.

6. Chiang, T.; Duncan, F.E.; Schindler, K.; Schultz, R.M.; Lampson, M.A. Evidence that weakened centromere cohesion is a leading cause of age-related aneuploidy in oocytes. Curr Biol 2010, 20, 1522–1528, doi:10.1016/j.cub.2010.06.069.

7. Li, C.J.; Lin, L.T.; Tsai, H.W.; Chern, C.U.; Wen, Z.H.; Wang, P.H.; Tsui, K.H. The Molecular Regulation in the Pathophysiology in Ovarian Aging. Aging Dis 2021, 12, 934–949, doi:10.14336/ad.2020.1113.

8. Webber, L.; Davies, M.; Anderson, R.; Bartlett, J.; Braat, D.; Cartwright, B.; Cifkova, R.; de Muinck Keizer-Schrama, S.; Hogervorst, E.; Janse, F.; et al. ESHRE Guideline: management of women with premature ovarian insufficiency. Hum Reprod 2016, 31, 926–937, doi:10.1093/humrep/dew027.

9. Coccia, M.E.; Rizzello, F. Ovarian reserve. Annals of the New York Academy of Sciences 2008, 1127, 27–30.

10. Zhang, J.-G.; Xu, C.; Zhang, L.; Zhu, W.; Shen, H.; Deng, H.-W. Identify gene expression pattern change at transcriptional and post-transcriptional levels. Transcription 2019, 10, 137–146.

11. Lackner, D.H.; Bähler, J. Translational control of gene expression: from transcripts to transcriptomes. International review of cell and molecular biology 2008, 271, 199–251.

12. Kan, R.L.; Chen, J.; Sallam, T. Crosstalk between epitranscriptomic and epigenetic mechanisms in gene regulation. Trends in Genetics 2022, 38, 182–193.

13. Minarovits, J.; Banati, F.; Szenthe, K.; Niller, H.H. Epigenetic regulation. Patho-Epigenetics of Infectious Disease 2016, 1–25.

14. Holliday, R. DNA methylation and epigenetic mechanisms. Cell biophysics 1989, 15, 15–20.

15. Marshall, K.L.; Rivera, R.M. The effects of superovulation and reproductive aging on the epigenome of the oocyte and embryo. Molecular reproduction and development 2018, 85, 90–105.

16. Moghadam, A.R.E.; Moghadam, M.T.; Hemadi, M.; Saki, G. Oocyte quality and aging. JBRA assisted reproduction 2022, 26, 105.

17. Todeschini, A.-L.; Georges, A.; Veitia, R.A. Transcription factors: specific DNA binding and specific gene regulation. Trends in genetics 2014, 30, 211–219.

18. Sirotkin, A.V. Transcription factors and ovarian functions. Journal of cellular physiology 2010, 225, 20–26.

19. Uhlenhaut, N.H.; Treier, M. Forkhead transcription factors in ovarian function. Reproduction 2011, 142, 489.

20. Glisovic, T.; Bachorik, J.L.; Yong, J.; Dreyfuss, G. RNA-binding proteins and post-transcriptional gene regulation. FEBS letters 2008, 582, 1977–1986.

21. Kishore, S.; Luber, S.; Zavolan, M. Deciphering the role of RNA-binding proteins in the post-transcriptional control of gene expression. Briefings in functional genomics 2010, 9, 391–404.

22. Zhao, B.S.; Roundtree, I.A.; He, C. Post-transcriptional gene regulation by mRNA modifications. Nature reviews Molecular cell biology 2017, 18, 31–42.

23. Solyga, M.; Majumdar, A.; Besse, F. Regulating translation in aging: from global to gene-specific mechanisms. EMBO Reports 2024, 25, 5265–5276.

24. Masuda, K.; Marasa, B.; Martindale, J.L.; Halushka, M.K.; Gorospe, M. Tissue-and age-dependent expression of RNA-binding proteins that influence mRNA turnover and translation. Aging 2009, 1, 681.

25. Will, C.L.; Lührmann, R. Spliceosome structure and function. Cold Spring Harbor perspectives in biology 2011, 3, a003707.

26. Li, J.; Lu, M.; Zhang, P.; Hou, E.; Li, T.; Liu, X.; Xu, X.; Wang, Z.; Fan, Y.; Zhen, X. Aberrant spliceosome expression and altered alternative splicing events correlate with maturation deficiency in human oocytes. Cell Cycle 2020, 19, 2182–2194.

27. Zhu, Z.; Xu, W.; Liu, L. Ovarian aging: mechanisms and intervention strategies. Medical Review 2023, 2, 590–610.

28. Lambert, S.A.; Jolma, A.; Campitelli, L.F.; Das, P.K.; Yin, Y.; Albu, M.; Chen, X.; Taipale, J.; Hughes, T.R.; Weirauch, M.T. The human transcription factors. Cell 2018, 172, 650–665.

29. Wang, J.; Shi, A.; Lyu, J. A comprehensive atlas of epigenetic regulators reveals tissue-specific epigenetic regulation patterns. Epigenetics 2023, 18, 2139067.

30. Liao, J.-Y.; Yang, B.; Zhang, Y.-C.; Wang, X.-J.; Ye, Y.; Peng, J.-W.; Yang, Z.-Z.; He, J.-H.; Zhang, Y.; Hu, K.; et al. EuRBPDB: a comprehensive resource for annotation, functional and oncological investigation of eukaryotic RNA binding proteins (RBPs). Nucleic Acids Research 2019, 48, D307–D313, doi:10.1093/nar/gkz823.

31. Chen, W.; Moore, M.J. Spliceosomes. Current Biology 2015, 25, R181–R183.

32. Park, S.U.; Walsh, L.; Berkowitz, K.M. Mechanisms of ovarian aging. Reproduction 2021, 162, R19–R33.

33. Londero, A.P.; Orsaria, M.; Tell, G.; Marzinotto, S.; Capodicasa, V.; Poletto, M.; Vascotto, C.; Sacco, C.; Mariuzzi, L. Expression and prognostic significance of APE1/Ref-1 and NPM1 proteins in high-grade ovarian serous cancer. American journal of clinical pathology 2014, 141, 404–414.

34. Brüning, A.; Blankenstein, T.; Jückstock, J.; Mylonas, I. Function and regulation of MTA1 and MTA3 in malignancies of the female reproductive system. Cancer and Metastasis Reviews 2014, 33, 943–951.

35. Zhou, C.; Li, D.; He, J.; Luo, T.; Liu, Y.; Xue, Y.; Huang, J.; Zheng, L.; Li, J. TRIM28-mediated excessive oxidative stress induces cellular senescence in granulosa cells and contributes to premature ovarian insufficiency in vitro and in vivo. Antioxidants 2024, 13, 308.

36. Zhu, Q.; Chen, J.; Pan, P.; Lin, F.; Zhang, X. UBE2N regulates paclitaxel sensitivity of ovarian cancer via Fos/P53 axis. OncoTargets and therapy 2020, 12751–12761.

37. Broekmans, F.; Soules, M.; Fauser, B. Ovarian aging: mechanisms and clinical consequences. Endocrine reviews 2009, 30, 465–493.

38. Wilkosz, P.; Greggains, G.D.; Tanbo, T.G.; Fedorcsak, P. Female reproductive decline is determined by remaining ovarian reserve and age. PloS one 2014, 9, e108343.

39. Hickey, M.; Elliott, J.; Davison, S.L. Hormone replacement therapy. Bmj 2012, 344.

40. Wu, C.; Chen, D.; Stout, M.B.; Wu, M.; Wang, S. Hallmarks of ovarian aging. Trends in Endocrinology & Metabolism 2025.

41. López-Otín, C.; Blasco, M.A.; Partridge, L.; Serrano, M.; Kroemer, G. The hallmarks of aging. Cell 2013, 153, 1194–1217.

42. Charames, G.S.; Bapat, B. Genomic instability and cancer. Current molecular medicine 2003, 3, 589–596.

43. Muraki, K.; Nyhan, K.; Han, L.; Murnane, J.P. Mechanisms of telomere loss and their consequences for chromosome instability. Frontiers in oncology 2012, 2, 135.

44. Abdul, Q.A.; Yu, B.P.; Chung, H.Y.; Jung, H.A.; Choi, J.S. Epigenetic modifications of gene expression by lifestyle and environment. Archives of pharmacal research 2017, 40, 1219–1237.

45. Kaushik, S.; Cuervo, A.M. Proteostasis and aging. Nature medicine 2015, 21, 1406–1415.

46. Trifunovic, A.; Larsson, N.G. Mitochondrial dysfunction as a cause of ageing. Journal of internal medicine 2008, 263, 167–178.

47. Van Deursen, J.M. The role of senescent cells in ageing. Nature 2014, 509, 439–446.

48. Ruzankina, Y.; Brown, E. Relationships between stem cell exhaustion, tumour suppression and ageing. British journal of cancer 2007, 97, 1189–1193.

49. Li, L.; Shi, X.; Shi, Y.; Wang, Z. The signaling pathways involved in ovarian follicle development. Frontiers in Physiology 2021, 12, 730196.

50. Baruzzo, G.; Hayer, K.E.; Kim, E.J.; Di Camillo, B.; FitzGerald, G.A.; Grant, G.R. Simulation-based comprehensive benchmarking of RNA-seq aligners. Nature methods 2017, 14, 135–139.

51. Morris, D.R.; Geballe, A.P. Upstream open reading frames as regulators of mRNA translation. Molecular and cellular biology 2000, 20, 8635–8642.

52. Vo, K.; Sharma, Y.; Paul, A.; Mohamadi, R.; Mohamadi, A.; Fields, P.E.; Rumi, M.K. Importance of transcript variants in transcriptome analyses. Cells 2024, 13, 1502.

53. Pant, P.; Chitme, H.; Sircar, R.; Prasad, R.; Prasad, H.O. Genome-wide association study for single nucleotide polymorphism associated with mural and cumulus granulosa cells of PCOS (polycystic ovary syndrome) and non-PCOS patients. Future Journal of Pharmaceutical Sciences 2023, 9, 27.

54. Hughes, C.H.; Smith, O.E.; Meinsohn, M.-C.; Brunelle, M.; Gévry, N.; Murphy, B.D. Steroidogenic factor 1 (SF-1; Nr5a1) regulates the formation of the ovarian reserve. Proceedings of the National Academy of Sciences 2023, 120, e2220849120.

55. Zamberlam, G.; Lapointe, E.; Abedini, A.; Rico, C.; Godin, P.; Paquet, M.; DeMayo, F.J.; Boerboom, D. SFRP4 is a negative regulator of ovarian follicle development and female fertility. Endocrinology 2019, 160, 1561–1572.

